# Tsetse blood-meal sources, endosymbionts, and trypanosome infections provide insight into African trypanosomiasis transmission in the Maasai Mara National Reserve, a wildlife-human-livestock interface

**DOI:** 10.1101/2020.04.06.027367

**Authors:** Edward Edmond Makhulu, Jandouwe Villinger, Vincent Owino Adunga, Maamun M. Jeneby, Edwin Murungi Kimathi, Enock Mararo, Joseph Wang’ang’a Oundo, Ali Abdulahi Musa, Lillian Wambua

## Abstract

**Background:** African trypanosomiasis (AT) is a neglected disease of both humans and animals caused by *Trypanosoma* parasites, which are transmitted by obligate hematophagous tsetse flies (*Glossina* spp.). Understanding of AT transmission is hampered by limited knowledge on interactions of tsetse flies with their vertebrate hosts and the influence of endosymbionts on vector competence, especially in wildlife-human-livestock interfaces. We identified the tsetse species, their blood-meal sources, and the correlation between endosymbiont and trypanosome infection status in the trypanosome-endemic Maasai Mara National Reserve (MMNR) of Kenya.

**Methodology/Principal Findings:** Among 1167 tsetse flies (1136 *Glossina pallidipes*, 31 *Glossina swynnertoni*) collected from 10 sampling sites, 28 (2.4%) were positive by PCR for trypanosomes, majority (17/28) being *Trypanosoma vivax*. Blood-meal analyses based on high-resolution melting analysis of mitochondrial cytochrome c oxidase 1 and cytochrome b gene PCR products (n = 345) identified humans as the most common vertebrate host (37%), followed by hippopotamus (29.1%), African buffalo (26.3%), elephant (3.39%), and giraffe (0.84%). Trypanosome-infected flies had fed on hippopotamus and buffalo. Additionally, PCR analysis revealed that tsetse flies were more likely to be infected with trypanosomes if they were infected with the *Sodalis glossinidius* endosymbiont (P = 0.0022 Fisher’s exact test).

**Conclusions/Significance:** Diverse species of wildlife hosts may contribute to the maintenance of tsetse populations and/or persistent circulation of African trypanosomes in the MMNR. Although the African buffalo is known to be a key reservoir of AT, the higher proportion of hippopotamus blood-meals in trypanosomes-infected flies identified here indicates that other wildlife species may also be important to transmission cycles. No trypanosomes associated with human disease were identified, but the high proportion of human blood-meals identified are indicative of human African trypanosomiasis transmission risk. Furthermore, this work provides data showing that *Sodalis* endosymbionts can is associated with increased trypanosome infection rates in endemic ecologies.

**Author summary:** Human and animal African trypanosomiasis are neglected tropical diseases with potential to spread to new areas. Wild animals are important reservoirs for African trypanosomes and crucial in the emergence and re-emergence of AT. Vertebrate host-vector-parasite interactions are integral to trypanosome transmission. We investigated the vertebrate blood-meals and trypanosomes-endosymbionts co-infections in tsetse flies, which have been associated with reservoirs and vector competence, respectively, on AT transmission in Kenya’s Maasai Mara National Reserve. We identified tsetse fly diversity, trypanosome and endosymbiont infection status, and vertebrate blood-meal hosts to infer potential transmission dynamics. We found that *Glossina pallidipes* was the major tsetse fly vector and that *Trypanosoma vivax* was the main trypanosome species circulating in the region. Humans, hippopotamus, and buffalo were the most frequented for blood-meals. Buffalo and hippopotamus blood-meals were identified in trypanosome infected flies. Feeding of the flies on both humans and wildlife may potentiate the risk of the human trypanosomiasis in this ecology. Additionally, we found that the endosymbiont *Sodalis glossinidius* is associated with higher trypanosome infection rates in wild tsetse flies. These findings emphasize the importance of understanding the interaction of tsetse flies with vertebrate blood-meal sources and their endosymbionts in the transmission and control of AT.

## Introduction

African trypanosomes (genus *Trypanosoma*), cyclically transmitted by the tsetse fly vector (genus *Glossina*), cause a group of diseases known as African trypanosomiasis (AT). In man, the disease is called sleeping sickness (human African trypanosomiasis, HAT), while in animals it is called nagana (African animal trypanosomiasis, AAT). African trypanosomiasis is endemic in 37 countries in Africa, in regions inhabited by the insect vector. Approximately 70 million people and 60 million cattle in AT endemic regions are at risk of infection [1,2]. Consequently, reduced productivity due to chronic disease in humans animal and loss of livestock through death, particularly in regions where pastoralism is the main economic activity threatens food security, quality of living, and economic stability [3,4]. Therefore, more effective control and management strategies of AT are required.

Control of AT has involved combinations of active surveillance, vector control strategies, and mass chemotherapy [5]. Notably, chemotherapy has been limited by increasing levels of resistance to the available trypanocides, chemotoxicity, and unavailability of new drugs [5,6]. To address limitations associated with chemotherapy, transmission disruption through vector control is crucial. Vector control is largely applied in areas where livestock are kept [7,8]. However, wild animals sustain the life cycles of tsetse flies [9,10] and the parasites they transmit [11,12], and are thus an important factor of transmission dynamics of AT, particularly in wildlife ecologies. Tsetse fly blood-meal sources are highly variable, especially in wildlife areas. Hence, one sampling area cannot be used to make a generalized conclusion of tsetse feeding behavior [12]. Consequently, identification of tsetse fly host blood-meal sources in specific regions can help elucidate potential wild animals involved in AT transmission and provide baseline for research towards improving vector-control strategies, particularly in wildlife-human-livestock interfaces that serve as hotspots for the emergence and re-emergence of AT.

Moreover, transmission of trypanosomes from an infected animal to another by tsetse flies is highly influenced by vector competence – the vector’s ability to successfully acquire and transmit a pathogen. Competence is influenced by various factors including genotype, sex, species, immune status, and endosymbionts, [13–17]. Endosymbionts have been shown to influence the susceptibility of tsetse flies to trypanosomes [18]. *Wigglesworthia glossinidia, Sodalis glossinidius*, and *Wolbachia pipentis* are well-defined tsetse fly endosymbionts with direct and indirect effects on the tsetse fly vectorial capacity [19]. Despite numerous studies on the influence of endosymbionts on vectorial competence [13,14,20–22], studies on the presence and influence of tsetse fly endosymbionts in wildlife-livestock-human interfaces are scant.

The Maasai Mara National Reserve (MMNR) is a prime tourist destination in Kenya that is surrounded by a number of ranches and is thus characterized by constant interactions between wildlife and humans and their livestock. With endemic tsetse fly populations, there have been recent cases of tourists contracting HAT in the MMNR [23,24]. Therefore, the MMNR is an ideal study site for investigating the contribution of tsetse fly blood-meal sources and the major endosymbionts of tsetse flies in relation to transmission of African trypanosomes in a human-livestock-wildlife interface. We sought to understand the interactions occurring among natural tsetse fly populations, trypanosome species, endosymbionts, and vertebrate hosts in the MMNR. Specifically, we investigated the diversity of tsetse fly species and their trypanosome species, vertebrate blood-meal sources, and *Sodalis, Wolbachia*, and salivary gland hypertrophy virus (SVGH) endosymbionts.

## Materials and methods

### Study area

Field sampling was carried out within the MMNR (1°29′24″S 35°8′38″E, 1500 m above sea level) located in southwest region of Kenya, which is contiguous with the Serengeti National Park (SNP) in Tanzania (Figure 1). This sampling site is located approximately 150 km south from the equator and covers an area of 1500 km2. The MMNR is home to a diverse variety of flora and fauna, and is famously known for its wild animals and ‘Great Migration’ of wildebeests, zebras, and antelopes across the Mara River. Grassland forms the major vegetation cover in this ecosystem, with swampy grounds found around the riverbanks. The sampling sites were selected along the rivers due to their high populations of animals (Figure 1). Ethical clearance for this research in protected areas was sought from and approved by the Kenya Wildlife Service (KWS) Research Authorization committee.

**Fig 1.**
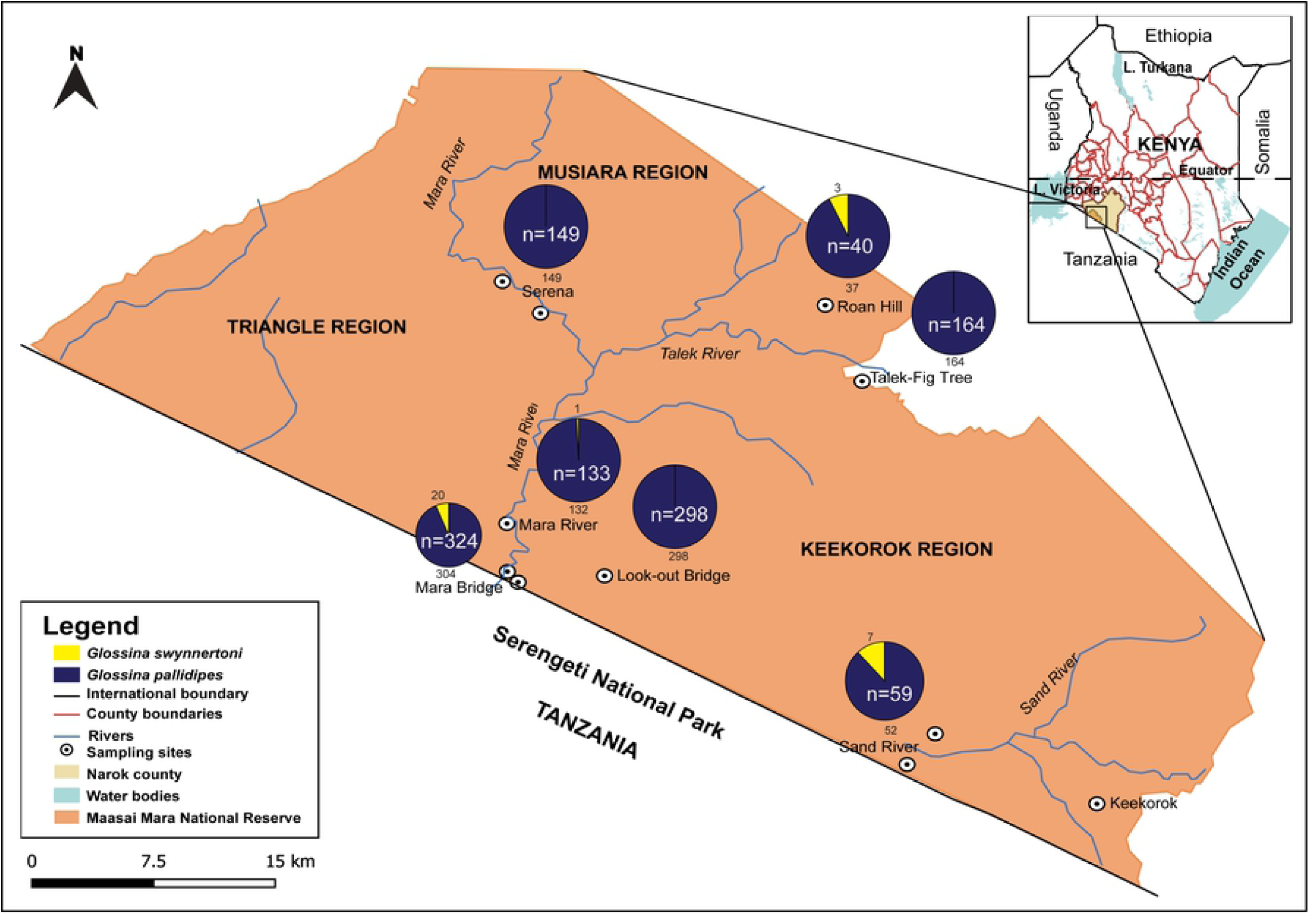
Map showing tsetse fly sampling sites from Maasai Mara National Reserve and number of tsetse species sampled. Each sampling site has its total sampled flies indicated in the pie chart.

### Tsetse collection and identification

Tsetse flies were trapped at the start of the annual wildebeest migration between June and July 2016 using Nguruman (Ngu) traps baited with acetone and cow urine. Traps were set in the morning (10 - 11 am) at different sampling sites in the various regions demarcated by Mara, Talek, and Sand Rivers, and at the wildlife crossing points across the Mara River at the border of Kenya and Tanzania’s SNP (Figure 1). The traps were emptied after 24 hours, and trapped flies were transferred into 50-mL falcon tubes and stored in dry ice before transportation in liquid nitrogen to the laboratory at the International Centre of Insect Physiology and Ecology (*icipe*), Nairobi where they were sorted. The flies were identified to species level under a light microscope (Stemi 2000-C, Zeiss, Oberkochen, Germany) based on standard published taxonomic keys [25].

### Nucleic acid extraction

Individual flies (after removal of legs and wings) were homogenized using six 2-mm zirconium beads in 1.5-ml microcentrifuge tubes using Mini-beadbeater-16 (BioSpecs Inc., Bartlesville, OK, USA) for 20 seconds. DNA was extracted from the homogenate of each sample using the ammonium acetate protein precipitation method described by Adams *et al*. [26], with slight modifications. Briefly, 300 µl of cell lysate buffer (10 mM Tris-HCl, pH 8.0, 0.5% SDS and 5 mM EDTA) was added into homogenized samples and incubated for 90 minutes at 65°C.

Thereafter, 100 µl of protein precipitate solution (8M ammonium acetate and 1M EDTA) was added to each mixture, which were vortexed for 30 seconds, incubated in ice for 30 minutes, and centrifuged at 14,000 x g for 15 minutes at 4°C. The supernatants were transferred into new 1.5-ml microcentrifuge tubes containing 300 µl of isopropanol, and mixed gently by inverting 100 times and centrifuging at 14,000 x g for 30 minutes. The supernatants were pipetted off and subsequently, 300 µl of ice-cold 70% molecular grade ethanol was added to each pellet, gently mixed by inversion and centrifuged at 14,000 x g for 30 minutes. Ethanol was pipetted off and the pellets were air-dried overnight. The DNA pellets were solubilized by adding 100 µl of PCR grade water and quantified using a NanoDrop™ 2000 Spectrophotometer (Thermo Scientific, NJ, USA). Concentrations were adjusted to 50 ng/µl using PCR grade water.

### PCR identification of African trypanosomes

Trypanosome infections in flies were detected using trypanosome-specific ITS1 CF and BR primers (S1 Table) as described by Njiru *et al*. [27]. *Trypanozoon* species were further resolved using species-specific primers (S1 Table). Briefly, glycosylphosphatidylinositol-phospholipase C polypeptide (GPI-PLC) and serum resistant-associated (SRA) species-specific primers were used to identify *T. brucei brucei* and *T. brucei rhodesiense*, respectively, by PCR [28]. *Trypanosoma congolense savannah* was identified according to Masiga *et al*. [29].

PCR amplification was carried out in final reaction volumes of 20 µl containing 10.4 µl of PCR grade water, 1× GeneScript PCR reaction buffer and 1.6 units of Green Taq DNA polymerase enzyme (GeneScript, New Jersey, USA), 1 µl (final concentration 0.5 µM) of each primer and 200 ng DNA template. The PCR reactions were performed in a SimpliAmp™ Thermal Cycler (Applied Biosystems, California, USA) programmed as follows; initial denaturation step at 94°C for 3 minutes followed by 30 cycles of denaturation at 94°C for 30 seconds, annealing at a temperature specific for each primer (S1 Table) for 30 seconds and extension at 72°C for 45 seconds and a final extension at 72°C for 7 minutes. PCR grade water was used as a negative control, in place of DNA template. DNA obtained from characterized and archived stocks of African trypanosome species were used as positive controls. The PCR products were size separated by ethidium stained agarose gel electrophoresis and viewed under UV light. Gel images were captured using Genoplex (VWR International GmbH, Darmstadt, Germany) and images processed using Adobe Photoshop (Adobe Photoshop CC).

### Host blood-meal identification

Blood-meal sources were determined by PCR coupled with high-resolution melting (HRM) analysis of vertebrate cytochrome c oxidase subunit I (COI) and cytochrome b (cyt b) mitochondrial genes as previously described [30–32]. We analyzed 760 flies, representing 65% of the sampled population, which included all engorged flies (n = 39) and 721 randomly selected non-engorged flies. Final concentrations of the PCR reactions were performed in 20 µl PCR reaction volumes, which included 4 ul of 5× Hot FIREPol EvaGreen HRM Mix (Solis BioDyne, Teaduspargi, Estonia), 0.5 µM of each primer, 250 ng of DNA template, and 10 µl of PCR grade water. The PCR cycling conditions included an initial denaturation at 95°C for 15 minutes followed by 35 cycles of denaturation at 95°C for 30 seconds, annealing at specific temperatures for COI and cyt b primers (S1 Table) for 30 seconds and elongation at 72°C for 30 seconds. This was followed by a final extension at 72°C for 7 minutes. Thereafter, HRM analysis of PCR products was conducted as described by [30–32]. HRM profiles were analysed using the Rotor-Gene Q software v2.1 with normalized regions between 76.0-78.0°C and 89.50-90.0°C. Amplicons representative of each unique HRM profile were purified using ExoSAP-IT kit (USB Corporation, Cleveland, Ohio, USA) according to the manufacturer’s instructions and sequenced at Macrogen (South Korea). The sequences were analysed and aligned using the MAFFT plugin in Geneious software version 11.1.4 [33]. Vertebrate species were confirmed by sequence alignments with Basic Local Alignment search tool (BLAST) [34] hits obtained from the GenBank database.

### PCR identification of *Sodalis glossinidius, Wolbachia*, and salivary gland hypertrophy virus

We screened 760 tsetse flies for their endosymbionts, *S. glossinidius, Wolbachia*, and salivary gland hypertrophy virus (SGHV). The PCR reactions were done in 20 µl reaction volumes using endosymbiont-specific primers (SI table) and similar concentrations of PCR components as those described above for host blood-meal identification. The PCR cycles steps were the same as those used in host blood-meal analysis with changes made only in the annealing temperatures, which were specific for the different primer pairs [35–37]. Positive controls for *Wolbachia* and *Sodalis* were isolated from *Aedes aegypti* and *Glossina pallidipes*, respectively, while a plasmid standard from a synthetic construct of the *P74* gene of SGHV from GenScript was used as a positive control. PCR-grade water used to reconstitute the extracted genomic DNA was used as a negative control. The amplified products were size separated in 2% (W/V) agarose gels. The amplicons were visualized, images captured, and processed as previously described.

### Statistical analysis

Data were processed and analyzed through descriptive statistical tools using RStudio at 95% confidence levels. Two-tailed t-tests were used to compare variations of tsetse fly species proportions between the sampling blocks and the mean differences of host blood-meals between the tsetse fly species. Fisher’s exact test was used to determine whether endosymbiont infections were correlated with trypanosome co-infections.

## Results

### Tsetse fly species identified

A total of 1167 tsetse flies were collected from the ten sampling sites, of which 1136 were *G. pallidipes* and 31 were *G. swynnertoni* (Table 1). The highest number of *G. swynnertoni* flies (87.1%; n = 27) were sampled from sites closer to the border between the MMNR and the SNP (i.e. Sand River and Mara Bridge sampling sites) as depicted in Fig 1. Female tsetse flies constituted 61% of the collected samples, whereas 39% were male. There was no statistical difference between the mean sex proportions of *G. pallidipes* (0.963±0.029) and *G. swynnertoni* (0.037±0.029) (t (4) = 31.54, p < 0.001).

**Table 1.**
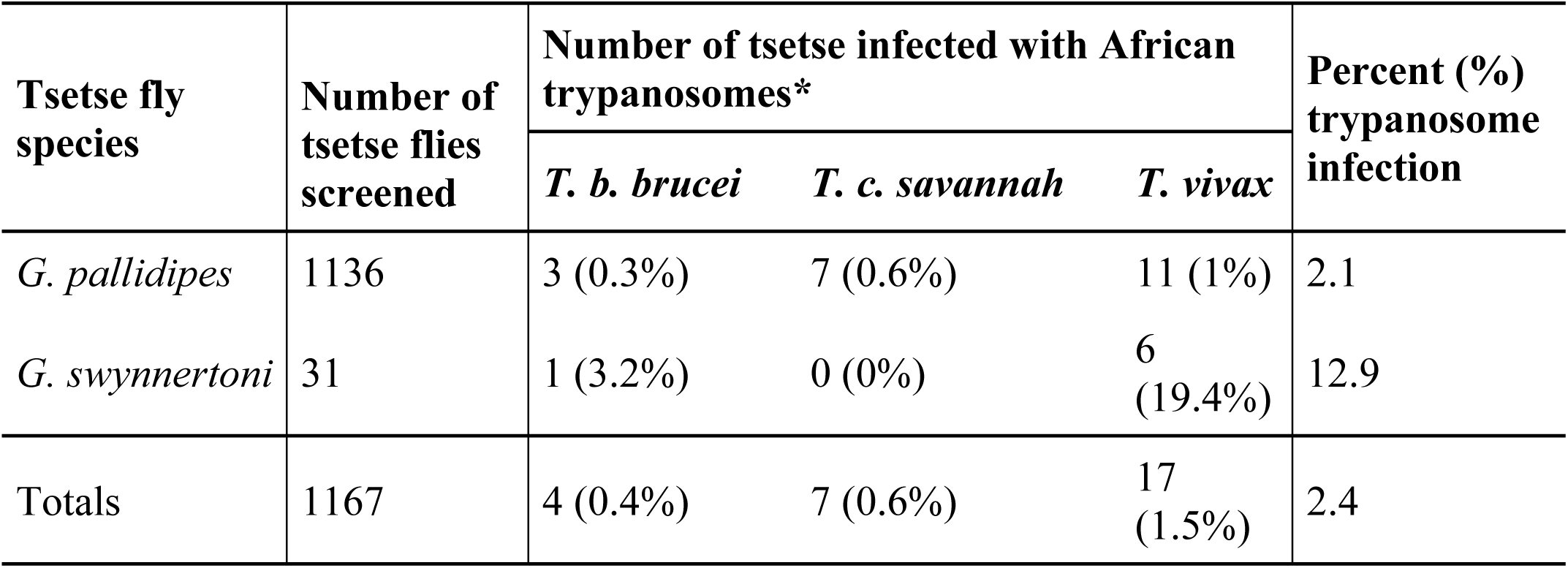
Numbers and proportions of trypanosome species identified in tsetse fly species. *The percentage of trypanosomes infecting each tsetse fly species is indicated in brackets.

### Trypanosome species identified in sampled tsetse flies

Trypanosome DNA was amplified in 28 (2.40%) out of the 1167 tsetse flies sampled (Table 1). The African trypanosome species identified were *T. vivax* (17/28), whereas 25% were *T. congolense savannah* (7/28), and 14.3% were *T. brucei brucei* (4/28) (S3 Table, GenBank accessions MK684364-MK684366). A representation of the samples positive for trypanosomes by PCR is shown in S1 Fig. Trypanosome infection rate was higher in *G. swynnertoni* (12.9%, n = 4/31) than in *G. pallidipes* (2.1%; n=24/1136). There were no mixed trypanosome infections in either of the fly species.

### Tsetse blood-meal sources identified

Vertebrate blood-meals were detected and identified in 46.6% (354/760) of the tsetse flies analyzed which comprised of 328 *G. pallidipes* and 26 *G. swynnertoni* (Fig. 2; S2 Table). The most common source of blood-meal was from humans (*Homo sapiens*) (n = 131) (S3 Table, GenBank accession MK684355, MK684357), followed by hippopotamus (*Hippopotamus amphibious*) (S3 Table, GenBank accession MK684356) (n = 103), African buffalo (*Syncerus caffer*) (S3 Table, GenBank accessions MK684354, MK684358) (n = 93), African savannah elephant (*Loxodonta africana*) (S3 Table GenBank accession MK684359) (n = 12), and giraffe (*Giraffa camelopardis*) (S3 Table, GenBank accession MK684360) (n = 3). There were 406 samples that had HRM peaks lower than 0.5 rate in fluorescence (dF/dT), thus qualified as having a non-detectable blood-meal.

**Fig 2.**
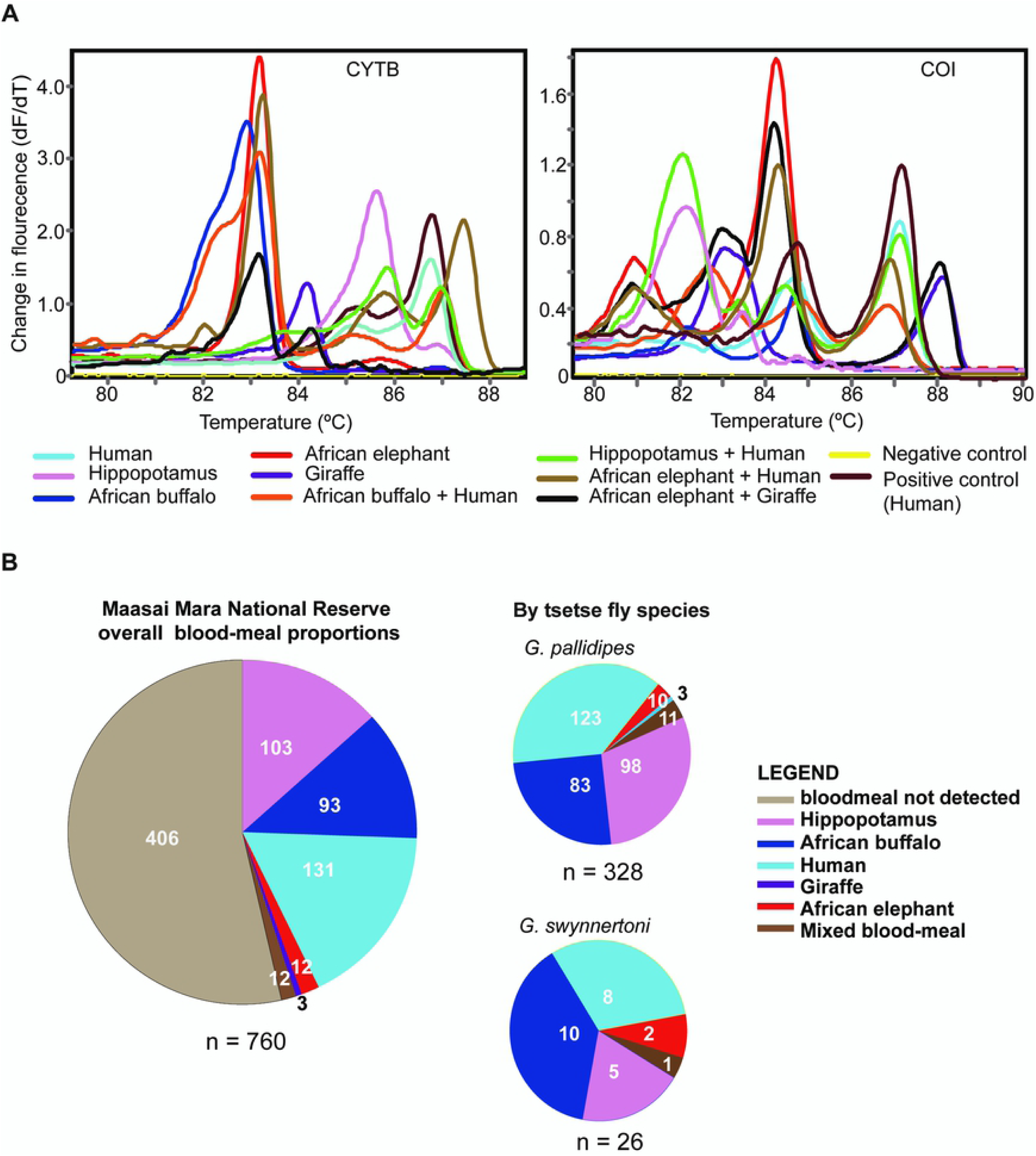
Blood-meal melt curves and proportions of vertebrate species identified. A. High resolution melt curves of single species and mixed species blood-meals. Mixed blood-meals were determined by matching melt profile peaks to those of more than one blood-meal control. B. Overall blood-meal proportions and proportions per tsetse species.

Humans were the most frequently identified blood-meal sources in *G. pallidipes*, whereas African buffalo was the major blood-meal sources for *G. swynnertoni* (Fig. 2B; S2 Table). However, there was no significant difference in the mean blood-meal source proportions between the two tsetse fly species (t (4) = 2.47, p = 0.069). We further observed that of the 28 trypanosome-infected tsetse, 14 (10 *G. pallidipes* and four *G. swynnertoni*) had blood-meals from African buffalo, while eight *G. pallidipes* had blood-meals from hippopotamus. The blood-meal sources for six trypanosome-infected flies could not be identified. The vertebrate blood-meal detection rates were 94.87% and 43.69% in engorged and non-engorged flies, respectively.

Twelve mixed blood-meals were detected (Fig 2), accounting for 3.4% of the positively identified samples. These samples had distinct melt curves that mapped onto those of known reference samples. Of these, human and buffalo mixed blood-meals was the most frequent combination (6/12), followed by human and elephant (3/12), elephant and giraffe (2/12), and human and hippopotamus (1/12) (Fig 2; S2 Table). Mixed blood-meals were further confirmed by sequencing of representative PCR-HRM amplicons (S2 Fig).

### Co-infection of tsetse with endosymbionts and African trypanosomes

A total of 69 (n = 760, 9.08%) flies (66 *G. pallidipes*, three *G. swynnertoni*) were infected with the endosymbiont *S. glossinidius* (S3 Table, 16S-23S rRNA GenBank accessions MK684361-MK684363) (S3 Fig.). Notably, a greater proportion of *S. glossinidius*-positive *G. pallidipes* flies were co-infected with trypanosomes (7/68, 10.3%) than *G. pallidipes* without *Sodalis* infections (16/661, 2.42%) (P = 0.0033 Fisher’s exact test) (Table 2). Five of the co-infections were with *T. congolense* and two were with *T. vivax*. As only one *G. swynnertoni* fly, which was also trypanosome infected (*T. vivax*), was positive for *S. glossinidius*, the sample size was too small for meaningful association analysis in this tsetse species. Nonetheless, across both species, the proportion of *S. glossinidius* infected flies that were co-infected with trypanosomes (8/69, 11.6%) was similarly significantly higher than in tsetse without the *S. glossinidius* endosymbiont (20/691, 2.89%) (P = 0.0022, Fisher’s exact test).

**Table 2.** Statistical analysis of *Sodalis glossinidus* and *Wolbachia* co-infection with trypanosomes in *G. pallidipes* and *G. swynnertoni*. Abbreviations: **S+/S-** *S. glossinidus* positive/negative, **W+/W-** *Wolbachia* positive/negative, **T+/T-** trypanosome positive/negative.

|  | *G. pallidipes* |  |  | *G. swynnertoni* |  |  | *G. pallidipes* |  |
| --- | --- | --- | --- | --- | --- | --- | --- | --- |
|  | T+ | T- |  | T+ | T- |  | T+ | T- |
| S+ | 7 (0.96%) | 61 (8.37%) | S+ | 1 (3.23%) | 0 (0.00%) | W+ | 1 (0.14%) | 17 (2.33%) |
| S- | 16 (2.19 %) | 645 (88.47%) | S- | 4 (12.90%) | 26 (83.87%) | W- | 22 (3.02%) | 689 (94.51%) |
| P = 0.0033*; $\alpha < 0.05$ | | | P = 0.1613; $\alpha > 0.05$ | | | P = 0.555; $\alpha > 0.05$ | | |

Eighteen flies (n = 760; 2.37%), 16 females and two males (all *G. pallidipes*), were infected with *Wolbachia* (S3 Table, GenBank accessions MK680053-MK680056) (S3 Fig). However, there was no statistical significance between *Wolbachia* and trypanosome co-infections in *G. pallidipes* (P = 0.555, Fisher’s exact test). No SGHV was detected in this study and therefore there were no co-infection studies.

## Discussion

Transmission of vector-borne diseases is dependent on the vector competence and the interactions between their vectors and vertebrate hosts that are reservoirs of the parasites [12,38,39]. This cross-sectional study focused on the transmission of AAT in the MMNR, a wildlife ecology in Kenya, with respect to the sources of tsetse fly blood-meals and the potential impact of three endosymbionts on their infection with animal trypanosomes. Our findings revealed that humans, hippopotamus, and African buffaloes were the most frequent blood-meal sources of tsetse flies in the MMNR. We also found that the endosymbiont, *S. glossinidius*, was positively correlated with trypanosome infection in wild-caught *G. pallidipes* tsetse flies in the MMNR, supporting the hypothesis that *Sodalis* potentiates AAT transmission in tsetse flies [40–42]. However, we found no correlation between *Wolbachia* and trypanosome infections, and no evidence of SGHV endosymbionts in the tsetse populations analyzed. These findings emphasize the importance of understanding the complete spectrum of interactions amongst vertebrates, tsetse fly vectors, endosymbionts, and trypanosome parasites, particularly in the context of wildlife-livestock-human interfaces where emergence and reemergence of AAT and other vector-borne diseases are reported.

*Glossina pallidipes* was the most abundant tsetse species sampled in the MMNR in this study, while *G. swynnertoni* was less abundant. This finding corroborates previous studies in which these two savannah tsetse species were found to be predominant in the Maasai Mara-Serengeti ecosystem of Kenya and Tanzania [43–45]. As both species are competent vectors of human and animal trypanosomes [46,47], their presence highlights the persistent risk of AAT and HAT in the MMNR. Despite their lower abundance relative to *G. pallidipes, G. swynnertoni* had higher rates of infection with trypanosomes, underpinning its critical role in transmission of African trypanosomes as demonstrated by previous studies [48]. However, unlike *G. pallidipes*, which is widely distributed across many habitats in Kenya, the geographical range of *G. swynnertoni* is limited to a narrow belt within the Maasai Mara-Serengeti ecosystem, which has resulted in the prioritization of this tsetse species as a target for elimination in East Africa [49]. Extensive efforts have been employed over the last four decades to reduce *G. swynnertoni* populations using various techniques as comprehensively reviewed by Nagagi and co-workers [49]. These have included spraying with both residual and non-residual insecticides, use of mechanical traps and baits with insecticide-impregnated traps or cloth targets, and insecticide-treated animals as live mobile targets. Coordinated studies are needed to evaluate their effect on tsetse populations and quantify their impact in East Africa.

Despite recent cases of HAT (caused by *T. b. rhodensiense*) being reported in East Africa [50], the trypanosome species identified in this study are only those responsible for causing trypanosomiasis in animals. Kenya is currently classified by the WHO as a country with diminished incidence of HAT (<10 cases in the last decade), with the recent cases being reported in 2012 in tourists returning from the MNNR [23,24]. Nevertheless, the persistent presence of *G. pallidipes* and *G. swynnertoni*, which are competent vectors of *T. b. rhodensiense*, coupled with the relatively higher incidences of HAT in neighboring Tanzania and Uganda and increased tourism, reinforces the need for coordinated surveillance and diagnosis in the MMNR and other HAT foci in eastern Africa. With reference to animal trypanosomes, this study identified *T. vivax* as the most prevalent species, followed by *T. congolense* and *T. brucei brucei*. Our findings are congruent with previous findings within the East African savannah [51,52]. The higher numbers of flies infected with *T. vivax* may be due to differences in development cycles in tsetse flies; *T. vivax* has all its development stages in the fly’s proboscis unlike *T. congolense* and *T. brucei*, which establish in the fly mid-gut where they are affected by low pH, proteases, and lectins [53,54]. Moreover, *T. vivax* usually achieves higher parasitemia in hosts than do *T. congolense* and *T. brucei*, further increasing its chances of being transmitted to tsetse flies during blood-feeding on infected hosts [54]. Infection rates were similar for both female and male tsetse flies, as found in another recent study conducted in Mtito Andei, Kenya [47].

The greater abundance of *G. pallidipes*, but higher trypanosome infection rate observed in *G. swynnertoni*, in the MMNR highlights the need for understanding the difference in susceptibility between the two tsetse species. Vector susceptibility of *G. pallidipes* to mid-gut trypanosomes has been shown to be lower compared to *G. morsitans morsitans* and *G. morsitans centralis* [55,56]. Further still, it has been shown that tsetse protection against trypanosome invasion is different for *G. pallidipes* and *G. morsitans morsitans* [56]. Similarly, field studies have shown *G. swynnertoni* to be more susceptible than *G. pallidipes* [46,49]. Given that *G. swynnertoni* is an important species in the Mara-Serengeti ecosystem, its potentially greater susceptibility to trypanosome infection needs further investigation to elucidate its relative role in trypanosome transmission relative to the more abundant sympatric *G. pallidipes*.

Blood feeding of tsetse fly populations in the wild is influenced by the composition of vertebrate host species in an area and how these species attract tsetse flies [12]. Our identification of animal trypanosome DNA in flies with hippopotamus and African buffalo blood-meals was not surprising as these vertebrates are known to be reservoirs for *T. vivax, T. congolense*, and *T. brucei* [12,55]. Nevertheless, these results suggest that active transmission cycle of animal trypanosomiasis in this wildlife-livestock interface may be maintained by multiple potential vertebrate hosts. Despite the abundance of wildebeest, zebra, and other antelopes when the study was conducted during the Great Migration season, no blood-meals from these hosts were detected in the tsetse flies. These findings are congruent with previous reports that *G. pallidipes* and *G. swynnertoni* exhibit significant specificity in host selection whereby wildebeest are not preferred blood-meal sources [57,58], and that zebra skin odors are repellant to *G. pallidipes* [59]. The influx of people in the MMNR due to high tourism activities in the Great Migration season, may partially explain why humans were frequent blood-meal sources. Identification of mixed blood-meals from humans and wildlife is indicative of the inherent risk of HAT transmission in the MMNR [11,12], even though *T. b. rhodensiense* was not detected in this study.

Visual cues and odors released by vertebrate hosts influence tsetse fly host choice and have been pivotal to the development of baited traps and targets for the control and management of tsetse fly populations, HAT, and AAT. A tsetse repellant formulation mimicking the odor of waterbuck (*Kobus ellipsiprymnus defassa*), a non-host animal, was recently developed and used as an innovative collar device to protect cattle from tsetse bites and AAT [60]. Visual cues have been extensively exploited in the development of improved traps – stationery and mobile targets impregnated with insecticides for riverine/”palpalis” [61–63] and savannah/”morsitans” [49,64] groups of tsetse. However, it has been noted that for the morsitans group of tsetse flies, including *G. pallidipes* and *G. swynnertoni*, host odors play a more significant role than visual cues as they strongly attract the tsetse flies across long ranges of up to 100 m [65]. Acetone and butanone odors obtained from cattle have long been used as attractants of choice in tsetse fly control [66]. However, other better tsetse fly attractants, such as 2-propanol, have been identified [67]. Our observed high rates of buffalo, hippopotamus, and human blood-meals imply that semiochemicals from these vertebrates may be possible candidates to advance research for novel host-derived cues in control of *G. pallidipes* and *G. swynnertoni*. Better understanding of the molecular mechanisms of tsetse fly olfaction associated with host selection can help to evaluate candidate host semiochemicals as potential attractants or repellants [67–69]. Knowledge of emergent repellant odors (such as those described from zebra and waterbuck) [59,60], coupled with new host attractants, present unique opportunities to further improve tsetse bait technology using “Push-Pull” and/or “Attract-and-Kill” approaches.

This study further highlights the sensitivity of HRM technique to accurately, reliably, rapidly, and reproducibly identify arthropod blood-meal hosts. We were able to identify blood-meals from wild-caught non-engorged flies. Unlike serological and other PCR-based techniques for blood-meal identification [57,70,71], the use of HRM to detect sequence variants is fast, cost-effective, accurate, easy-to-use, and sensitive, making it a more economical tool for blood-meal analysis [30,72] and pathogen detection/identification [71,73,74]. Sequencing of representative samples with combined human-hippopotamus and human-African buffalo peaks confirmed their mixed blood-meal status (S2 Fig.).

Our finding that higher proportions of tsetse flies infected with the endosymbiont *S. glossinidus* were infected with trypanosomes than those without *S. glossinidius* corroborates previous studies that suggest that *S. glossinidius* infection may have the potential to increase the ability of both wild caught [40,41] and lab reared [42,75] tsetse flies to acquire trypanosomes. Further, this finding implies that *S. glossinidius* infection in tsetse populations may be used as a positive indicator of trypanosomiasis risk. Nonetheless, the functional role of *S. glossinidius* in tsetse flies was not explored in this study and knowledge on the same remains limited [76]. However, inhibition of tsetse mid-gut and mouthpart lectins by Nacetyl-d-glucosamine (GlcNAc), a product of chitin catabolism by *S. glossinidius*, has been proposed as the main factor associated with *S. glossinidius* and increased tsetse-vector competence [18,77,78]. Nevertheless, this association is complex and a number of other factors, including geographic location, tsetse fly species, sex, and age also affect the capacity of *S. glossinidius* to increase vector competence in wild-caught tsetse flies [77]. While more studies are needed to elucidate the role of *Sodalis* endosymbionts on tsetse competence to vector trypanosomes, our findings support that *S. glossinidius* infection increases probability of savannah tsetse flies to acquire animal trypanosome infection in this wildlife-livestock interface.

## Conclusions

Emergence and/or reemergence of AT, especially in human-wildlife-interfaces like the MMNR, where AT has been recently reported, happens occasionally. With limitations on current methods of control and management of AT and its tsetse fly vectors, more research on the factors influencing trypanosome transmission is required. Identification of trypanosome-infected tsetse flies that had fed on hippopotamus and African buffalo highlights these two vertebrate species as possible reservoirs of trypanosomes in the MMNR, providing a basis for investigating their contributions to AT in other human-wildlife-interfaces. Further understanding of the attractiveness of hippopotamus and expounding on existing knowledge on African buffalo to tsetse flies based on the volatiles they release may help improve tsetse baits and repellants. In addition, our findings demonstrate that the endosymbiont *S. glossinidius* increases tsetse fly susceptibility to trypanosome infection in this endemic ecology. These findings support the idea that *S. glossinidius* can be a potential target for vector control [79]. Despite *T. b. rhodensiense* not being detected, it has previously been reported in the MMNR. Hence, there is need for intense prevalence studies of *T. b. rhodensiense* in tsetse flies from the MMNR.

## Acknowledgments

The authors thank Daniel Ouso and Edwin Ogola (*icipe*) for their contributions in blood-meal analysis protocol and positive controls. Antoinette Miyunga, Stephen Mwiu, Vasco Nyaga, Dennis Lemayian and Richard Bolo (all of KWS) are acknowledged for their assistance in field sampling. We are also thankful to Mr. James Kabii (*icipe*) for his technical support and logistics in carrying out this project.

## Author’s contributions

Conceived and designed the experiments: LW, JV, MJ, VOA. Performed the experiments: EEM, EM, JWO, AAM. Analyzed the data: EEM, JV, LW, VOA, EMK. Tsetse collection: LW, JV, MJ. Tsetse fly identification: EEM. Wrote the paper: EEM, LW, JV, VOA, MJ, EMK. Manuscript revision: All authors.

## Supporting Information

**S1 Fig. PCR detection of trypanosome species in tsetse flies. A**. Agarose gel electrophoresis images of representative tsetse fly samples positive for Trypanozoon, *T. vivax*, and *T. congolense* PCR amplicons with ITS BR/CR primers specific for African trypanosomes species. The trypanozoon group were further resolved using primer pairs specific for *T. b. rhodesiense* and *T. b. brucei*. **B**. PCR amplification results for detection of *T. b. rhodesiense* using primers targeting the SRA gene. **C**. PCR amplification results for detection of *T. b. brucei* using primers targeting the TbbGPI-PLC gene. M represents the molecular ladder – represents negative control and + represents positive control.

**S2 Fig. DNA sequence analysis of mixed blood-meals. A**. Hippopotamus and human mixed blood-meal cytochrome b sequences aligned and edited using Geneious v8.0.1. **B**. Buffalo and human mixed blood-meal cyt b sequences aligned and edited using Geneious v8.0.1. Scientific names and the GenBank accession numbers highlighted in red represent sequences obtained from this study.

**S3 Fig. PCR detection of endosymbionts in tsetse flies. A**. Agarose gel electrophoresis image of a representative PCR amplicons of *S. glossinidus* DNA. **B**. Agarose gel electrophoresis image of a representative PCR amplicons for *Wolbachia*.

**S1 Table. Primer list with annealing temperatures**. Details of primer sequences and PCR conditions used in this study.

**S2 Table. Trypanosome species and host blood-meals among the tsetse fly species in this study**. Distribution of trypanosome infections and blood-meals sources in *Glossina pallidipes* and *Glossina swynnertoni* in Maasai Mara, Kenya.

**S3 Table. Data URL repository associated with this study**. Details of the nucleotide sequences generated in this study and URLs for obtaining the respective accessions in GenBank.

